# Functional export of NDM-7 to outer membrane vesicles in *Klebsiella pneumoniae* compromises imipenem and cefiderocol activity

**DOI:** 10.64898/2026.06.26.734736

**Authors:** Ana M. González, Valeria Quiroz, Katherine Soto, Christina M.A.P. Schuh, Lorena Diaz, César A. Arias, Alejandro J. Vila, José M. Munita, Carolina López

## Abstract

The carbapenemases KPC and NDM are the most widespread determinants of carbapenem resistance in *Klebsiella pneumoniae*. Whereas KPC is a soluble periplasmic serine-β-lactamase, NDM is a membrane-anchored metallo-β-lactamase (MBL), a feature that promotes its incorporation into outer membrane vesicles (OMVs). OMVs are naturally released nanoparticles that deliver diverse bioactive cargo, including enzymes, virulence factors, and signaling molecules, and may contribute to antibiotic resistance. Here, we investigated the export and activity of carbapenemases in OMVs produced by carbapenem-resistant *Klebsiella pneumoniae* clinical isolates expressing NDM-7, an emerging variant, or KPC-2, as well as in isogenic laboratory-derived *K. pneumoniae* strains producing NDM-1, NDM-7 or KPC-2. NDM enzymes were detected in vesicles released by NDM-producing strains, whereas KPC-2 remained confined to the cellular fraction and was not observed in OMVs. OMVs contained catalytically active NDM enzyme and conferred protection to susceptible *K. pneumoniae* against imipenem. Importantly, NDM-positive OMVs also partially restored bacterial growth in the presence of cefiderocol, a siderophore cephalosporin used to treat infections caused by MBL producers. This protective effect was more pronounced for NDM-7 than for NDM-1. Together, these findings show that the clinically emerging NDM-7 variant is efficiently packaged into OMVs in *K. pneumoniae* and remains enzymatically active, allowing extracellular antibiotic degradation and conferring protection to susceptible bacteria exposed to carbapenems and cefiderocol.

## INTRODUCTION

Carbapenem-resistant *Klebsiella pneumoniae* (CRKP) has emerged as a major global challenge. Infections due to CRKP are associated with elevated mortality and pose significant therapeutic challenges for clinicians worldwide (1–4). As a consequence, CRKP rank amongst the top critical priority pathogens according to the World Health Organization report (5).

Resistance to carbapenems in CRKP mainly arises from the production of carbapenemase enzymes (6). The serine-β-lactamase (SBL) KPC and the metallo-β-lactamase (MBL) NDM are the most widely disseminated carbapenemases in *K. pneumoniae* (7). Infections caused by KPC-producing strains can be potentially treated with several β-lactam-β-lactamase inhibitor (BL-BLI) combinations such as ceftazidime-avibactam, meropenem-vaborbactam and imipenem-relebactam, although resistance to these combinations has been reported (8,9). Management of MBL-producing strains is even more challenging. At the moment, the only commercially available option to treat MBL producers is aztreonam-avibactam. The FDA has recently approved cefepime-zidebactam, and cefepime-taniborbactam is in advanced clinical trials. In general, new treatments are not readily available in developing countries where the prevalence of MBL-producing pathogens is high. Moreover, resistance to taniborbactam has already been reported in some MBL variants, including NDM-9 and members of the IMP family (10). Cefiderocol, a novel siderophore cephalosporin with *in vitro* activity against MBL producers, has emerged as a promising therapeutic option to manage NDM-producing Enterobacterales (11,12). However, resistance to cefiderocol and clinical failures have been reported among NDM-producing isolates (13,14). This phenomenon is particularly concerning because NDM enzymes are unique among clinically relevant MBLs in retaining hydrolytic activity against cefiderocol (15,16). Consequently, treatment options for infections caused by NDM-producing pathogens remain extremely limited, leaving clinicians with only a handful of therapeutic alternatives.

NDMs are lipoproteins anchored to the inner leaflet of the outer bacterial membrane (11). Membrane anchoring enhances NDM’s stability and resistance to proteolysis under zinc limiting conditions, as those elicited at the infection sites (11). Previous studies also showed that membrane anchoring facilitates incorporation of NDM-1 into outer membrane vesicles (OMVs) (11,12,17–19). OMVs are 20 to 300 nm spherical bilayer liposomes, naturally released by Gram-negative bacteria during growth. Increasing evidence suggests that OMVs act as vehicles for the export of different bacterial products, including the transfer of genetic material (20–22). NDM incorporation into OMVs could increase enzyme availability at infection sites, providing protection against otherwise susceptible surrounding bacterial populations (11,19). Secretion of KPC (a soluble periplasmic protein) into OMVs has also been reported in clinical isolates of *K*. *pneumoniae* (23). Soluble MBLs can be selectively exported into OMVs depending on their interaction with the outer membrane (18), but evidence for KPC enrichment in *K. pneumoniae* OMVs remains inconclusive.

To date, more than 90 NDM variants have been described (24), most differing from NDM-1 by only 1 to 3 amino acid substitutions (25,26). While NDM-1 and NDM-5 predominate globally (27), the emergence of the rare NDM-7 variant has been increasingly reported in regions such as France (28), Italy (29) and Chile (30–32). NDM-7 harbors two mutations on the protein surface (D130N, M154L) outside the active site, accounting for a similar hydrolytic activity to NDM-1 (33). However, these changes increase the stability of NDM-7 in the bacterial periplasm upon zinc deprivation as compared to NDM-1 (34,35), potentially enhancing its stability within OMVs.

Here, we studied the export and activity of carbapenemases in OMVs produced by clinical invasive carbapenemase-producing CRKP expressing NDM-7 or KPC-2. To further dissect the contribution of individual carbapenemases, we complemented these analyses using isogenic *K. pneumoniae* laboratory strains expressing NDM-1, NDM-7 or KPC-2. We found that NDM enzymes were readily incorporated into OMVs produced by both clinical isolates and isogenic strains, whereas KPC-2 remained largely confined to the cellular fraction and was not detected in OMVs. Importantly, NDM-containing OMVs protected susceptible bacteria against imipenem and cefiderocol, highlighting a potential mechanism by which the emerging NDM-7 variant may extend β-lactam hydrolysis beyond the producing cell and contribute to antimicrobial resistance against last generation compounds in *K. pneumoniae*.

## RESULTS AND DISCUSSION

### Clinical CRKP isolates show preferential export of active NDM-7 into OMVs

To investigate OMV production among clinical carbapenemase-producing CRKP, we selected five isolates recovered from bloodstream infections representative of dominant Chilean sequence types: ST11, ST25, ST45, ST15 (all harboring *bla*_NDM-7_) and ST1161, which carried *bla*_KPC-2_ (Table 1). Strains exhibited resistance to all carbapenems, while remaining susceptible to aztreonam–avibactam and cefiderocol (Table 1). Cefiderocol MICs among *bla*_NDM-7_-harboring isolates were higher (2–4 µg/mL) than in the *bla*_KPC-2_-producing isolate (0.25 µg/mL), with one NDM-7 strain exhibiting a MIC close to the resistance breakpoint. While all isolates remained susceptible to cefiderocol, the elevated MICs observed among NDM-7 producers are consistent with previous reports describing reduced cefiderocol susceptibility and the emergence of resistance in NDM-producing isolates (36–38).

**Table 1.** Characteristics and antimicrobial susceptibility profiles of clinical *Klebsiella pneumoniae* isolates.

| ID | ST | Isolation Year | Carbapenemase gene | Other β-lactamase genes | MIC (µg/mL) |  |  |  |  |  |
| --- | --- | --- | --- | --- | --- | --- | --- | --- | --- | --- |
|  |  |  |  |  | ERT | MEM | IPM | ATM | ATM/AVI | FDC |
| SCL 13085 | 11 | 2022 | *bla*<sub>NDM-7</sub> | *bla*<sub>SHV-11</sub> | ≥128 | ≥128 | 64 | 0.25 | 0.25 | 4 |
| SCL15912 | 25 | 2022 | *bla*<sub>NDM-7</sub> | *bla*<sub>CTX-M-2</sub>, *bla*<sub>SHV-11</sub>,<br>*bla*<sub>TEM-1</sub> | ≥128 | ≥128 | ≥128 | ≥64 | 0.5 | 2 |
| SCL 13788 | 45 | 2022 | *bla*<sub>NDM-7</sub> | *bla*<sub>CTX-M-15</sub>, *bla*<sub>SHV-1</sub>,<br>*bla*<sub>TEM-1</sub> | ≥128 | ≥128 | ≥128 | ≥64 | 0.25 | 2 |
| SCL 14690 | 15 | 2022 | *bla*<sub>NDM-7</sub> | *bla*<sub>CTX-M-15</sub>, *bla*<sub>SHV-28</sub>,<br>*bla*<sub>TEM-1</sub> | ≥128 | ≥128 | ≥128 | ≥64 | 0.125 | 2 |
| SCL 4611 | 1161 | 2019 | *bla*<sub>KPC-2</sub> | *bla*<sub>CTX-M-2</sub>, *bla*<sub>CTX-M-3</sub>,<br>*bla*<sub>SHV-187</sub>, *bla*<sub>TEM-1</sub> | ≥128 | 64 | 32 | ≥64 | 0.5 | 0.25 |

MIC, minimal inhibitory concentration; ERT, ertapenem; MEM, meropenem; IPM, imipenem; ATM, aztreonam; AVI,
avibactam; FDC, cefiderocol.

We purified OMVs from all five clinical isolates, as described in the experimental section. TEM images of the vesicle fraction revealed the presence of spherical, bilayered structures consistent with OMV production in all samples (Figure 1A). The total OMV yield was similar across the clinical isolates (Figure 1B). Nanoparticle tracking analysis (NTA) showed a size distribution ranging from ∼160 to 230 nm (Figure 1C), in agreement with the expected size range for OMVs (21).

**Figure 1.**
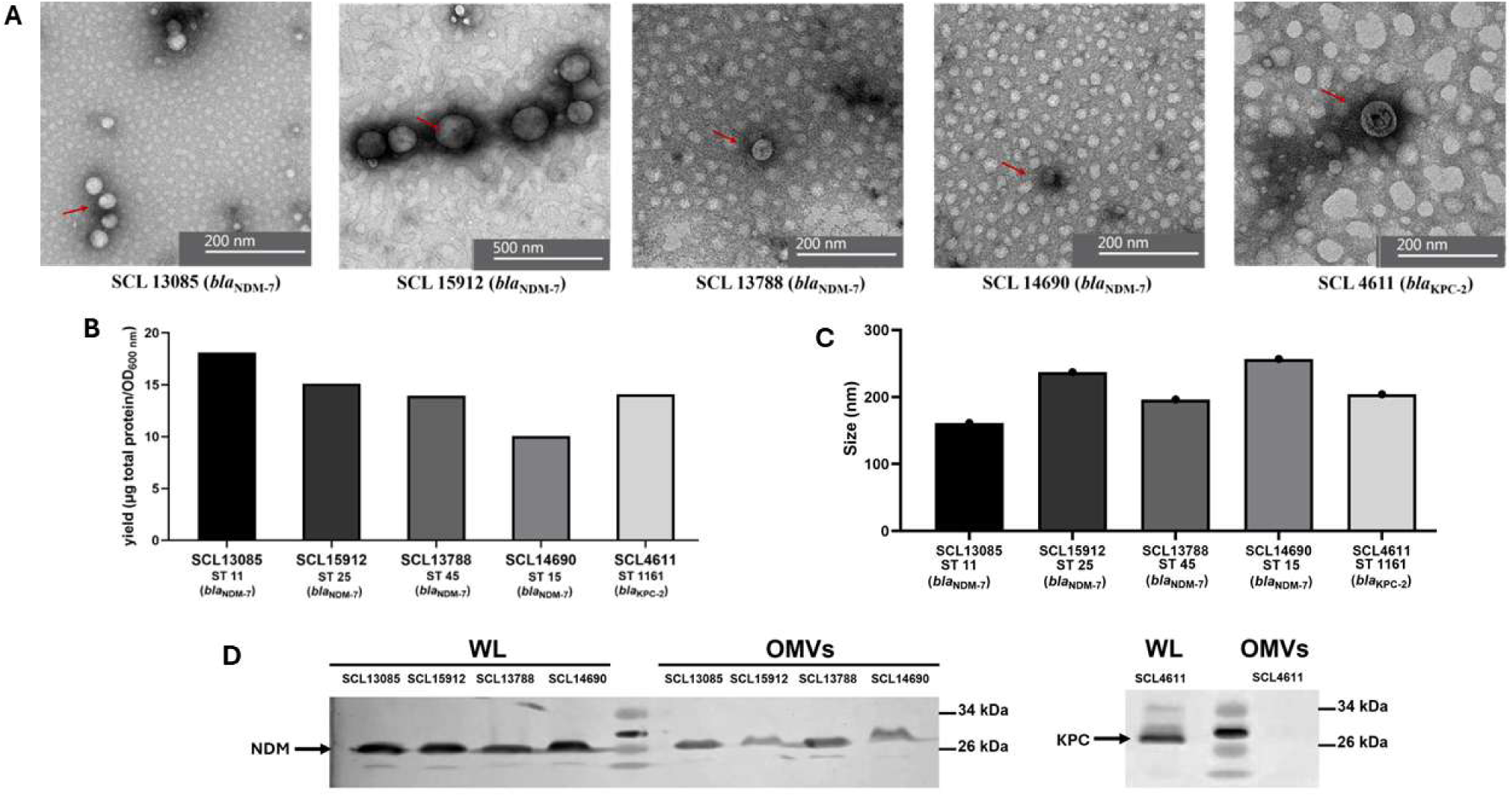
Purification and characterization of outer membrane vesicles (OMVs) from different CRKP sequence types (ST). (A) Transmission electron microscopy images of the OMVs isolated from CRKP strains. The red arrow highlights the presence of vesicles in the sample. (B) OMV yields obtained from bacteria based on total protein content from three independent purifications. (C) Mode of the OMVs diameter in CRKP strains from three independent experiments using NanoParticle Tracking Analysis (NTA). (D) NDM (∼28 kDa) or KPC (∼28.5 kDa) presence in whole lysates (WL) and outer membrane vesicles (OMVs).

Western blot assays in whole-cell lysates confirmed high levels of NDM-7 and KPC-2 enzymes in *bla*_NDM-7_- and *bla*_KPC-2_-harboring strains, respectively. However, while NDM-7 was readily detected in OMVs derived from all *bla*_NDM-7_-producers, KPC-2 was not detected in the vesicle fraction obtained from the isolate harboring *bla*_KPC-2_ (Figure 1D). These findings indicate that NDM-7 is efficiently incorporated into OMVs released by clinical CRKP isolates. Since OMVs can act as a mechanism for releasing misfolded, inactive proteins (39), we then evaluated the β-lactamase activity of isolated vesicles using qualitative nitrocefin hydrolysis assays (Figure S1). OMVs derived from all *bla*_NDM-7_ isolates produced a visible color change from yellow to red, consistent with β-lactam hydrolysis, indicating NDM-7 is not only selectively exported into OMVs, but it is present in a catalytically active form. A faint color change was observed with OMVs harvested from the *bla*_KPC-2_-positive isolate, suggesting the presence of β-lactamase activity despite the absence of detectable KPC-2 by western blot. Whole-genome sequencing (WGS) revealed the presence of additional SBL genes in this isolate including *bla*_CTX-M_, *bla*_SHV_ and *bla*_TEM_ genes (Table 1), which may contribute to the activity observed. However, as these genes were identified by WGS of the bacterial isolate, the presence of the corresponding enzymes in OMVs remains to be experimentally demonstrated.

To further dissect the contribution of different β-lactamase classes to the observed OMV-associated hydrolysis, nitrocefin assays were subsequently performed in the presence of class-specific inhibitors. EDTA, a metal-chelating agent that inactivates MBLs by removing Zn(II) from the active site, and avibactam, a non-β-lactam inhibitor active against KPC-2 and other SBLs enzymes were used. In OMVs derived from *bla*_NDM-7_-positive isolates, EDTA markedly reduced or abolished nitrocefin hydrolysis, whereas avibactam had little or no detectable effect. Conversely, avibactam abolished the nitrocefin color change observed in OMVs derived from the *bla*_KPC-2_-positive isolate, whereas EDTA did not have any observable effect (Figure S1). These findings support that the residual OMV-associated nitrocefin hydrolysis likely corresponds to the activity of additional SBLs encoded by this strain.

Notably, inhibitor responses varied among OMVs derived from blaNDM-7-positive isolates (Figure S1). EDTA almost completely abolished nitrocefin hydrolysis in OMVs isolated from SCL13085 and SCL14690, whereas a weaker inhibitory effect was observed for OMVs derived from SCL15912 and SCL13788. In SCL15912-derived OMVs, avibactam reduced hydrolytic activity more effectively than EDTA, suggesting the contribution of SBLs in addition to NDM-7. In contrast, the slight residual hydrolysis detected in OMVs from SCL13788 following EDTA treatment was not affected by avibactam and may reflect incomplete inhibition of OMV-associated NDM-7 under the endpoint assay conditions. Collectively, these data indicate clinical CRKP isolates produce OMVs carrying catalytically active β-lactamases, which was predominantly associated with the presence of NDM-7 (Figure 1 and Figure S1).

### Selective β-lactamase cargo in OMVs from isogenic *K. pneumoniae* strains

To better explore the export of carbapenemases into OMVs under controlled conditions, we decided to express different β-lactamases in an isogenic background, thereby eliminating differences among clinical strains. We transformed *K. pneumoniae* ATCC 13883 strains with pMBLe plasmids encoding *bla*_NDM-1_, *bla*_NDM-7_, or *bla*_KPC-2_. Antimicrobial susceptibility profiles of the isogenic strains are shown in Table S1. As expected, strains carrying *bla*_NDM_ alleles exhibited markedly increased MICs to all carbapenems as compared to the parental strain and the empty-vector control. The *bla*_KPC-2_ transformant also exhibited higher β-lactam MICs, although to a lesser extent. Aztreonam MICs remained unchanged in NDM producing strains, consistent with the lack of hydrolytic activity of MBLs against monobactams. Aztreonam-avibactam MICs were similarly preserved across all constructs. Cefiderocol MICs were higher in (2 µg/mL) as compared to the parental strain and the empty vector control (0.25 µg/mL). Despite this increase, all strains remained susceptible to cefiderocol. Western blot assays confirmed that both NDM-1 and NDM-7 were successfully expressed and selectively exported into OMVs (Figure 2A). In contrast, while readily detected in whole-cell lysates, KPC-2 was undetectable in the vesicular fraction, in line with our observations in clinical CRKP isolates, further suggesting an efficient packaging of NDM enzymes into OMVs as compared to KPC.

**Figure 2.**
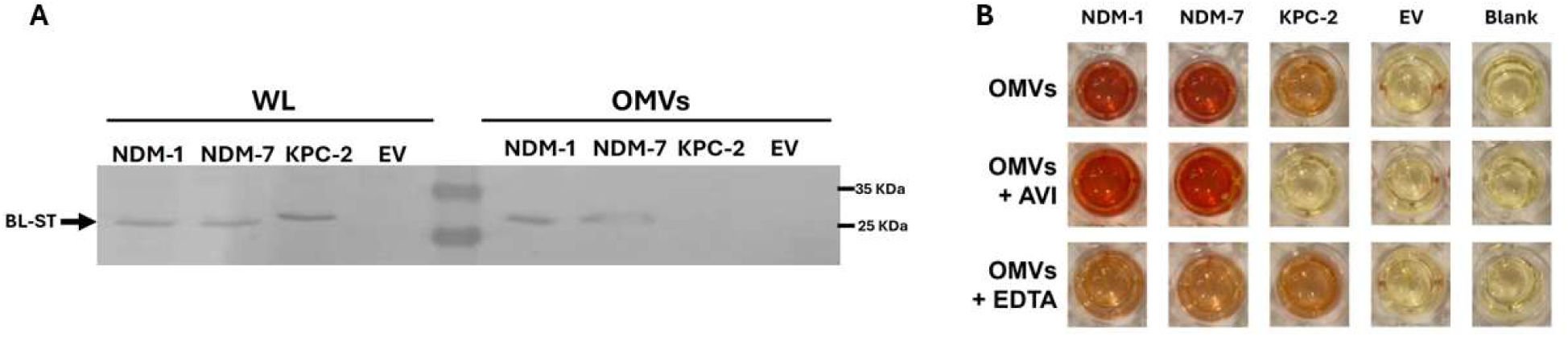
Isogenic *K. pneumoniae* strains reveal allele-specific carbapenemase – dependent OMV export and β-lactamase activity. (A) Western blot analysis of whole-cell lysates (WL) and purified outer membrane vesicles (OMVs) from *K. pneumoniae* ATCC 13883 isogenic strains expressing *bla*_NDM-1_, *bla*_NDM-7_, *bla*_KPC-2_, or the empty pMBLe vector (EV). (B) Nitrocefin hydrolysis assay using purified OMVs, measured in 96-well plates at 480 nm, based on the color change of nitrocefin from yellow to red. WL: whole lysate; AVI: avibactam; EDTA: ethylenediaminetetraacetic acid (metal chelating agent); BL-ST: Strep-tagged β-lactamase.

Consistent with these findings, functional analysis using a nitrocefin hydrolysis assays revealed strong β-lactamase activity in NDM-1 and NDM-7-containing OMVs. In this isogenic background, free of additional serine β-lactamase enzymes, β-lactamase activity of NDM-packed OMVs was completely abolished by EDTA and remained unaffected by avibactam, unambiguously demonstrating that the activity observed in clinical isolates was mediated by NDM rather than by serine β-lactamases (Figure S1 versus Figure 2B).

Taken together, our results support a differential incorporation of carbapenemases into OMVs. Both in clinical CRKP isolates and in isogenic *K. pneumoniae* strains, NDM carbapenemases (NDM-1 and NDM-7) were predominantly incorporated into OMVs, in contrast to KPC enzymes. The membrane-anchored nature of NDMs likely facilitates its selective packaging into OMVs, a phenomenon previously reported in *E. coli*, *Acinetobacter* spp. and *Pseudomonas* spp. (17). In contrast, KPC is a soluble periplasmic protein, a condition that may limit its incorporation into OMVs. Together, our findings highlight fundamental differences between NDM and KPC in their association with OMVs, with potential impact in their role facilitating antimicrobial resistance.

### NDM-loaded OMVs confer protection against imipenem

To evaluate the potential biological role of carbapenemase packaging into vesicles, we tested whether OMVs obtained from carbapenemase-producing clinical CRKP isolates could protect a susceptible control strain (*K. pneumoniae* ATCC 13883) against imipenem. Growth curve assays were performed with increasing imipenem concentrations and a fixed concentration of OMVs. OMVs derived from NDM-producing isolates conferred a clear protective effect, enabling growth of the ATCC strain at imipenem concentrations exceeding the clinical breakpoint (Figure 3B). In contrast, OMVs obtained from the KPC-producing isolate failed to confer protection, consistent with the absence of the enzyme export observed above.

**Figure 3.**
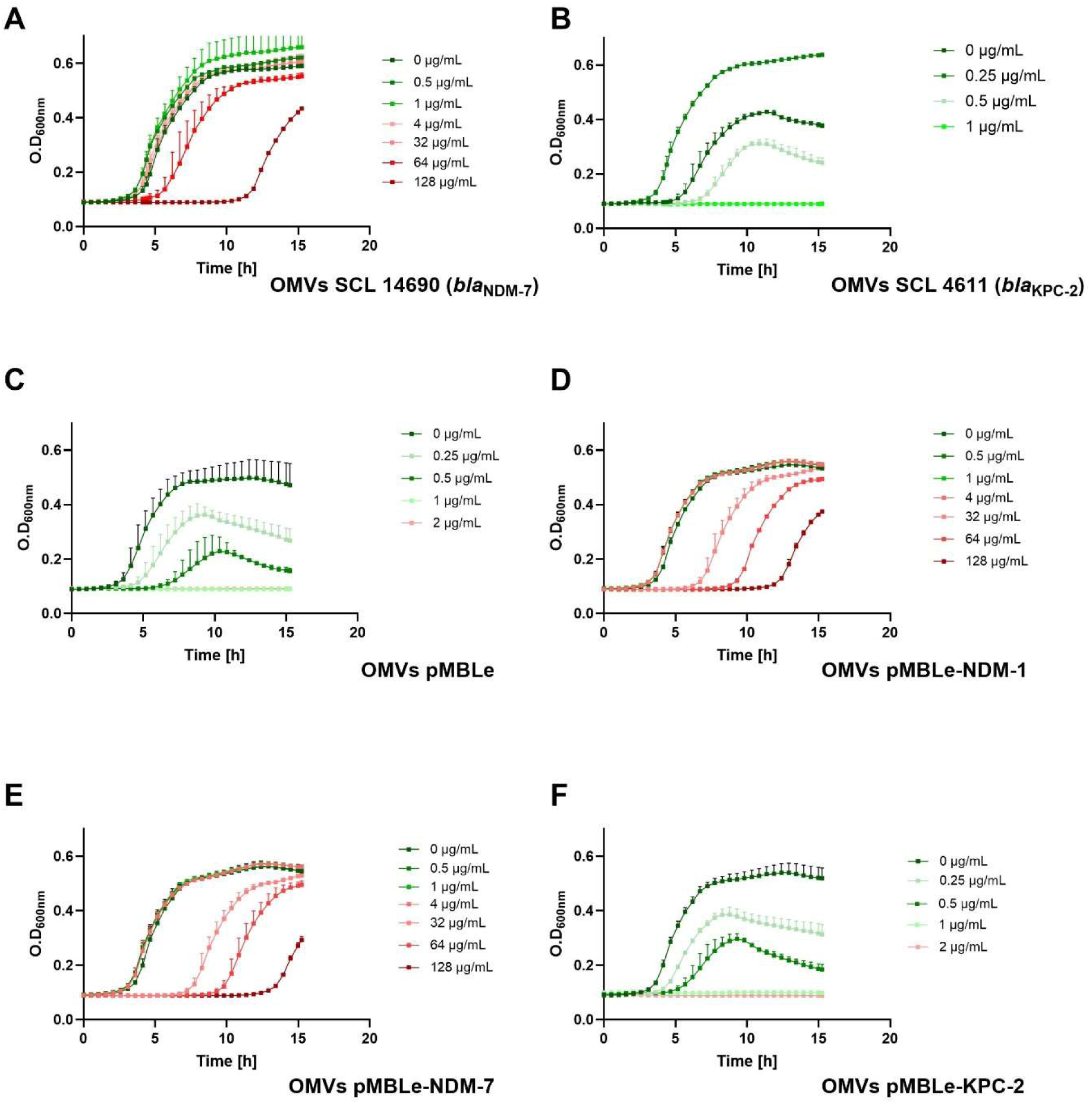
Representative growth curves of imipenem-susceptible ATCC 13883 in the presence of increasing concentrations of imipenem supplemented with 10 µg/mL OMVs purified from (A) SCL14690 (*bla*_NDM-7_), (B) SCL4611 (*bla*_KPC-2_), (C) pMBLe (empty vector), (D) pMBLe-NDM-1, (E) pMBLe-NDM-7, and (F) pMBLe-KPC-2. The established breakpoints for imipenem against Enterobacterales are ≤1 μg/mL for susceptibility (green) and >1 μg/mL for non-susceptibility (red).

These observations were further corroborated using our isogenic strain set, where OMVs from the *bla*_NDM7_-expressing construct conferred a similar protection against imipenem, while OMVs derived from *bla*_KPC-2_ or control strains exerted no effect (Figure 3C). Thus, both clinical and laboratory strains secrete NDM-packed OMVs loaded with carbapenemase activity and able to shield susceptible bacteria from the action of imipenem.

### NDM-positive OMVs attenuate susceptibility to cefiderocol

To determine whether OMVs from CRKP strains could modulate susceptibility to newer antimicrobials, we evaluated their protective effect against cefiderocol. Using the *K. pneumoniae* ATCC 13883 strain, growth assays were performed with increasing concentrations of cefiderocol, co-incubated with OMVs from selected CRKP isolates.

OMVs derived from an NDM-7-producing clinical strain partially restored bacterial growth at cefiderocol concentrations above the CLSI-defined susceptibility breakpoint (Figure 4A). These results suggest that the protective effect of OMVs packed with NDM extends to cefiderocol. In contrast, OMVs from the KPC-2-producing strain failed to protect susceptible bacteria from cefiderocol exposure. (Figure 4B).

**Figure 4.**
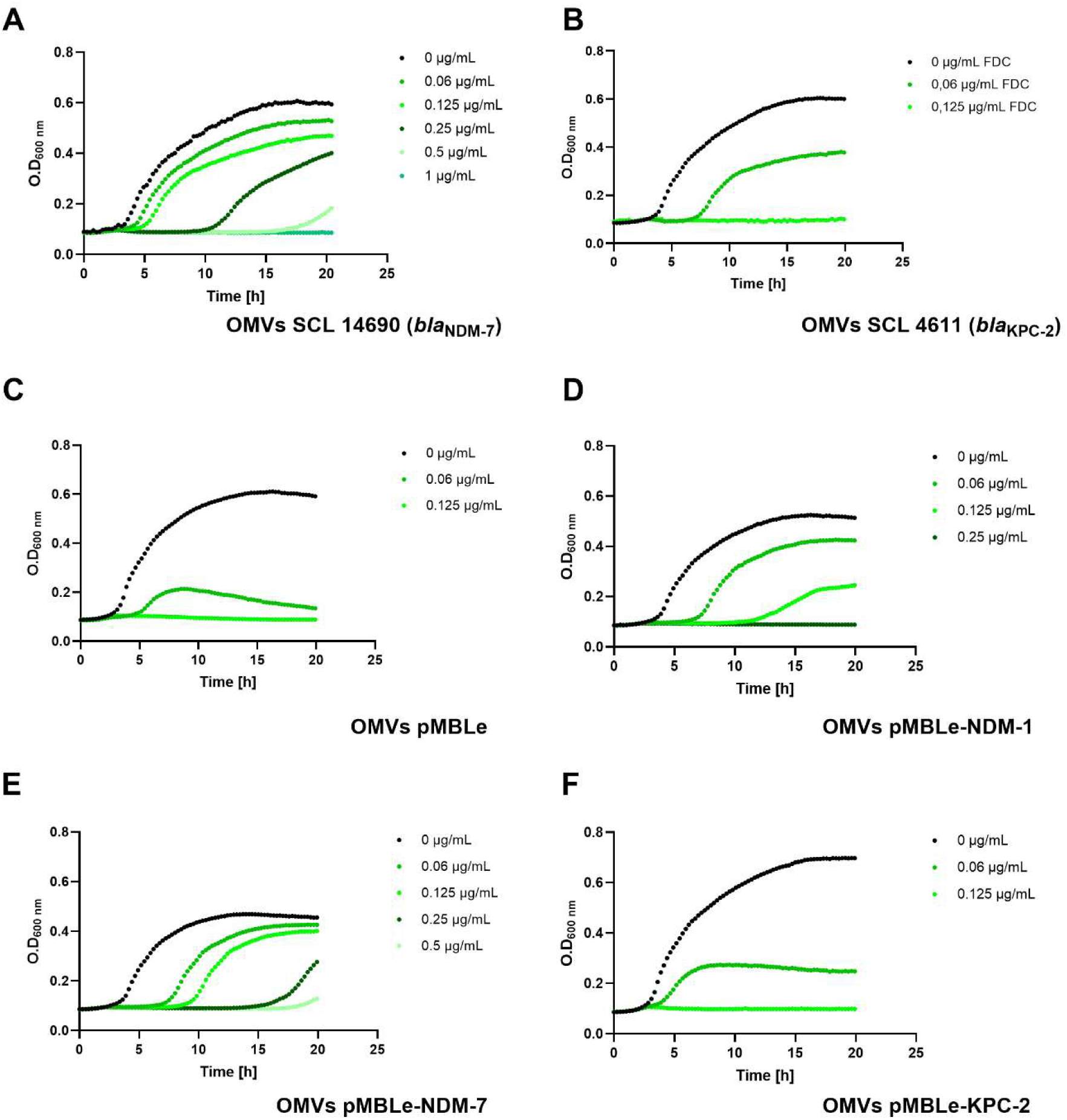
Representative growth curves of cefiderocol-susceptible ATCC 13883 in the presence of increasing concentrations of cefiderocol and 10 µg/mL OMVs purified from (A) SCL 14690 (*bla*_NDM-7_), (B) SCL 4611 (*bla*_KPC-2_), (C) pMBLe (empty vector), (D) pMBLe-NDM-1, (E) pMBLe-NDM-7, and (F) pMBLe-KPC-2. Cefiderocol (FDC) was tested at the indicated concentrations. The established breakpoints for FDC against Enterobacterales are ≤4 μg/mL for susceptibility (green) and >4 μg/mL for non-susceptibility (red).

To understand whether the NDM protecting effect against cefiderocol was variant-specific, we extended the analysis to the isogenic strains harboring *bla*_NDM-1_ and *bla*_NDM-7_. OMVs from the NDM-7-producing strain (pMBLe-NDM-7) similarly enabled growth of the ATCC strain at cefiderocol concentrations above 4 µg/mL (Figure 4D). In contrast, OMVs obtained from the NDM-1-expressing strain produced a weaker effect (Figure 4D). OMVs from cells expressing KPC-2 and those with the empty vector failed to confer any protection against cefiderocol (Figures 4C and 4F).

Our results indicate that OMVs carrying NDM carbapenemases reduce the activity of cefiderocol in the extracellular environment, possibly by NDM-mediated hydrolysis. While the degree of protection did not exceed clinical breakpoints, these findings highlight a potential vesicle-mediated mechanism that may contribute to the emergence of cefiderocol resistance under selective pressure. Additionally, under these experimental conditions, our results suggest NDM-7 may display higher protective effects than NDM-1.

Overall, we demonstrate that NDM-7 is efficiently packaged into OMVs released by invasive CRKP isolates and that exported NDM-7 retained enzymatic activity, supporting the role of OMVs as vehicles for the extracellular dissemination of carbapenemase activity. In contrast to previous observations, we did not observe efficient KPC-2 export into OMVs (23,40). This difference could be due to differences on bacterial backgrounds or induction conditions. In line with this variability, proteomic studies in OMVs have detected carbapenemase genes such as *bla*_KPC_ or *bla*_NDM_ in vesicle-associated DNA, while the corresponding proteins were not consistently identified (41,42), suggesting that their incorporation may occur at variable levels. Previous studies have also shown that membrane-associated carbapenemases, including NDM and OXA variants, are efficiently incorporated into OMVs, whereas soluble carbapenemases display more variable levels of vesicular export (18,43). Although our data do not directly address the mechanisms underlying KPC exclusion from OMVs, they further support the notion that carbapenemase incorporation into vesicles is a selective process influenced by both enzyme-specific and bacterial host-related factors. Indeed, envelope stress and detoxification responses have been proposed to modulate OMV cargo composition and promote the export of periplasmic proteins under specific conditions (17). Therefore, the factors determining carbapenemase export into vesicles deserve further consideration.

Selective packaging of NDM into OMVs may therefore enhance the extracellular distribution of active carbapenemase within bacterial communities. Notably, NDM enzymes are the only clinically-relevant MBL family capable of hydrolyzing cefiderocol, one of the few available options against these infections (13). In this context, OMV packaging of active NDM-7 may attenuate cefiderocol activity in the extracellular environment, potentially contributing to treatment failure and facilitating the persistence and dissemination of these worrisome pathogens. The potential impact of this mechanism may be especially relevant in regions where NDM-producing CRKP are highly prevalent, including many low- and middle-income countries, where these pathogens represent a major therapeutic challenge.

## MATERIALS AND METHODS

### Bacterial strains

Six carbapenemase-producing CRKP isolates recovered from clinical samples between 2019 and 2023 were used to evaluate the export of carbapenemase enzymes in OMVs. A complete genomic and phenotypic characterization of these clinical isolates is provided in Table 1. In addition, *K. pneumoniae* ATCC 13883 was used as an isogenic background strain and independently transformed with the pMBLe plasmid carrying different carbapenemase genes: *bla*_NDM-1_, *bla*_NDM-7_, and *bla*_KPC-2_, as previously described (11). Briefly, strains were grown under aerobic conditions at 37°C in Luria-Bertani (LB) broth, Mueller Hinton medium or on LB agar plates, as appropriate. For pMBLe-transformed strains, the growth medium was supplemented with gentamicin (30 µg/mL). Carbapenemase expression was induced with 20 µM IPTG. The strain harboring the empty pMBLe vector was used as a carbapenemase-negative control.

### Antimicrobial susceptibility testing

Minimum inhibitory concentrations (MIC) to ertapenem (ETP), meropenem (MEM), imipenem, aztreonam (ATM), aztreonam/avibactam (ATM/AVI) and cefiderocol were determined using broth microdilution according to CLSI recommendations (44). Measurements were performed in triplicate; quality control strains *E. coli* ATCC 25922 and *P. aeruginosa* ATCC 27853 were included in all assays.

### OMV isolation

OMVs were isolated as previously described (45). Briefly, clinical CRKP isolates were grown on LB agar and single colonies were selected and cultured in 5 mL LB broth at 37°C with constant shaking to an OD_600_ =1. Aliquots of 200 µL were then spread onto LB agar plates (n=15 plates per strain). In the case of isogenic strains derived from *K. pneumoniae* ATCC 13883 harboring different pMBLe constructs, agar plates were supplemented with 20 µM IPTG and gentamicin 30 µg/mL. Plates were incubated overnight at 37°C until confluent growth was achieved. Cells were recovered, resuspended in 35 mL of sterile PBS 1X, and centrifuged at 5000x g for 30 min at 4°C. Pellets were discarded, and supernatants were collected and centrifuged again under the same conditions to remove residual cell debris. The resulting supernatants were filtered through 0.22 μm pore-size filters and concentrated to a final volume of 200 µL using 100 kDa Amicon® Ultra Centrifugal Filter (Millipore). To verify sterility, 10 μL of each filtered supernatant was plated onto LB agar plates and incubated overnight at 37°C. OMVs preparations were stored at −80 °C until use.

### Characterization of OMVs

OMVs obtained from both clinical CRKP isolates and isogenic transformants were all characterized using *i)* transmission electron microscopy (TEM), *ii)* protein and lipid quantification assays, and *iii)* nanoparticle tracking analysis (NTA).

For transmission electron microscopy (TEM), the OMV suspension was diluted 1:100, and 10 µL of the diluted sample was deposited onto Formvar/carbon-coated 300-mesh copper grids (Ted Pella Inc., Redding, CA, USA). The grids were negatively stained with uranyl acetate and allowed to air-dry. TEM images were acquired using a Talos F200C G2 scanning transmission electron microscope (Thermo Fisher Scientific) operated at 200 kV at the UMA-UC Advanced Microscopy Center, Pontificia Universidad Católica de Chile, Santiago, Chile.

Protein concentration was measured using the Qubit Protein BR Assay Kit in a Qubit 4 Fluorometer (Thermo Fisher Scientific) according to manufacturer’s instructions. Lipid content was assessed using the FM 4-64™ dye (Thermo Fisher Scientific). Briefly, 5 μL of purified OMVs was mixed with 8 μL of 50 μg/mL FM 4-64 dye and PBS was added to a total volume of 200 μl in a Nunc 96-well culture plate (Thermo Fisher Scientific). After incubation at 37°C for 10 minutes, the plate was read using an Infinite® 200PRO M PLEX microplate reader with an excitation wavelength of 485 nm, an emission wavelength of 620 nm and gain of 143. PBS was used as a negative control.

Particle concentration and size distribution were determined by NTA using a Nanosight NS300 instrument (Malvern Instruments). OMV preparations were diluted 1:200 in sterile filtered PBS. Three 30 s videos were captured per sample (camera level = 11), processed (detection threshold = 5) and analyzed to determine size distribution and quantity of particles (software version: NTA 3.4 Build 3.4.4; Malvern Instruments Ltd).

### Carbapenemase detection and evaluation of β-lactamase activity

The presence of carbapenemase enzymes in OMVs was assessed by SDS-PAGE followed by Western blotting. For NDM-producing strains, a primary anti-NDM antibody was used at a 1:1,000 dilution from a 700 μg/mL solution (Instituto de Salud y Ambiente del Litoral, ISAL, UNL-CONICET, Argentina), followed by a goat anti-rabbit IgG (H+L) secondary antibody at a 1:5,000 dilution from a 1 mg/mL stock (Invitrogen). KPC detection was performed using anti- KPC antibody at a final concentration of 1 μg/mL, followed by a goat anti-mouse IgG (H+L) secondary antibody at a 1:5,000 dilution from 1 mg/mL stock. In the case of isogenic strains derived from *K. pneumoniae* ATCC 13883 transformed with the pMBLe constructs, a monoclonal anti-Strep Tag antibody was used at a 1:1,000 dilution from a 1 mg/mL stock (Sigma), followed by a goat anti-mouse IgG (H+L) secondary antibody at a 1:5,000 dilution from a 1 mg/mL stock. Briefly, samples were mixed with loading buffer and heated for denaturation. Proteins were separated by 10% SDS-PAGE and subsequently transferred onto a nitrocellulose membrane using a Trans-Blot® Turbo™ Transfer System (Biorad). OMV samples were normalized based on total protein and lipid content. Colorimetric detection of alkaline phosphatase activity was performed using 1-Step^TM^ NBT/BCIP (Thermo Fisher Scientific).

β-lactamase activity of carbapenemases exported in purified OMVs was assessed using a nitrocefin-based hydrolysis assay. Briefly, 10 μg of OMV from each isolate was incubated in 96 well-microplates with 10 μL of a 1 mg/mL nitrocefin solution in a final volume of 150 μL. Reactions were carried out at 37°C. As a negative control, OMVs were replaced with an equivalent volume of 1X PBS. To assess enzyme class specificity, parallel reactions were performed in the presence of 20 mM EDTA or 20 µg/mL avibactam (AVI).

### Protection assays and growth curves

The potential protective effect of OMVs isolated from carbapenemase-producing CRKP isolates on the growth of a susceptible *K. pneumoniae* ATCC 13883 strain was evaluated by performing growth curves in the presence or absence of imipenem or cefiderocol. Briefly, an overnight culture of *K. pneumoniae* ATCC 13883 was diluted 1:100 in LB broth and dispensed into sterile 96-well microplates (Falcon) containing increasing concentrations of imipenem (range 0-128 µg/mL). Bacterial growth was monitored by measuring optical density at 600 nm every 30 min using an Infinite® 200PRO M PLEX microplate reader (Tecan) for 20 h at 37°C. Experiments assessing cefiderocol activity were performed using the same methodology, except that bacteria were grown in cation-adjusted Mueller-Hinton broth.

Negative control experiments for both imipenem and cefiderocol including wells containing OMVs in the absence of the susceptible strain, and growth controls without antimicrobials were included in every experiment.

## ACKNOWLEDGEMENTS

This work was supported by the REPARA Network Grant (Redes Federales de Alto Impacto, Subsecretaría de Ciencia y Tecnología de la Nación) to A.J.V.; by the Agencia Nacional de Investigación y Desarrollo (ANID, Chile) through FONDECYT Regular Grant No.1250588 to J.M.M. and FONDECYT Regular Grant No.1220803 to CS; by the Centros de Investigación y Desarrollo de Excelencia de Interés Nacional (SENTINET) Grant No. CIN250062 to J.M.M. and L.D.; and by an ANID National Doctoral Fellowship (Beca Doctorado Nacional No. 21220961) awarded to A.M.G. C.L. and A.J.V. are staff members of CONICET, Argentina. The authors are grateful to Viviana Villalba (IBR-CONICET) for her valuable technical assistance and to the Advanced Microscopy Unit (UMA-UC) for support with TEM imaging.

---

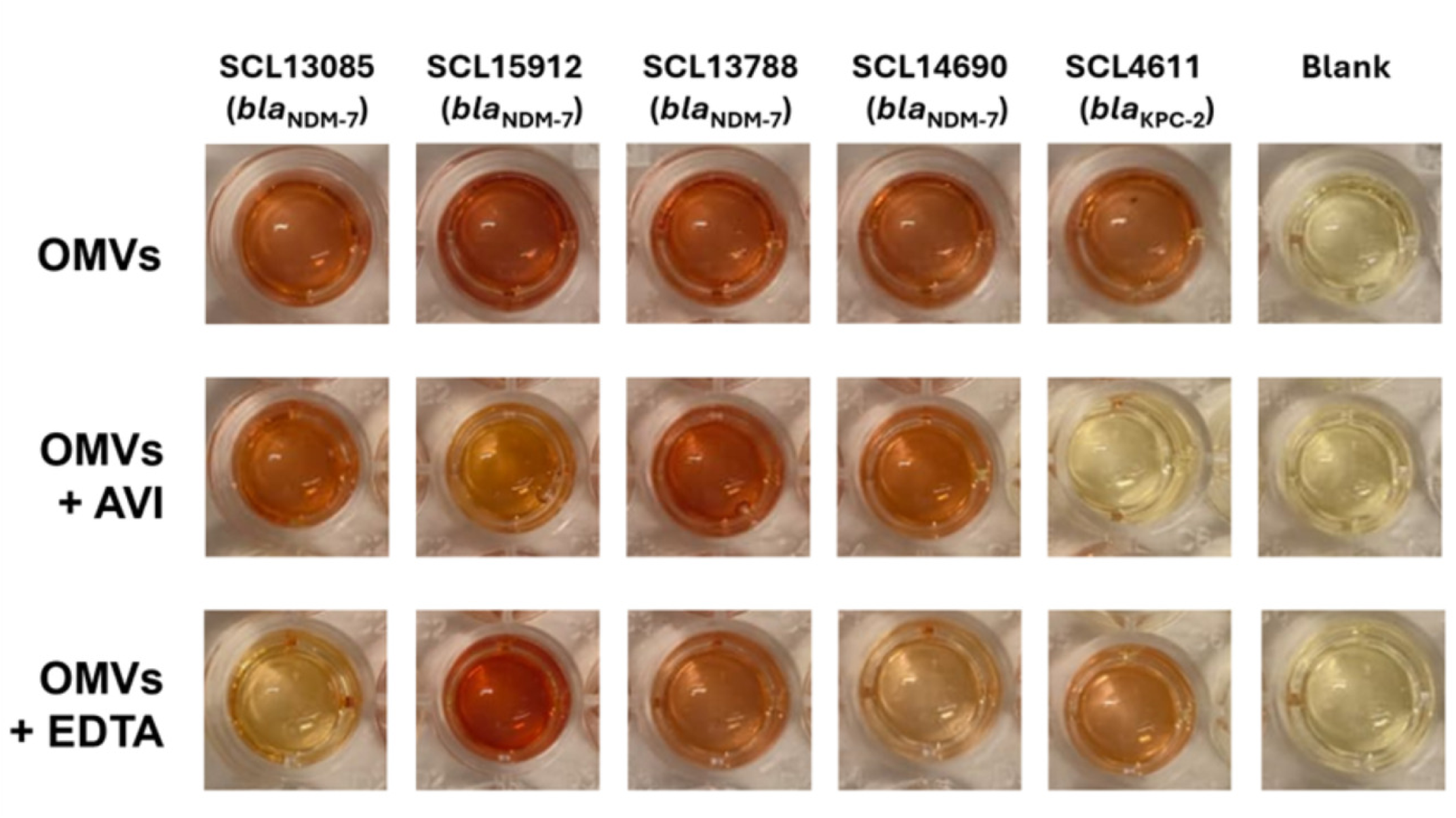

